# Cytoophidia safeguard binucleation of *Drosophila* male accessory gland cells

**DOI:** 10.1101/2022.09.27.509809

**Authors:** Dong-Dong You, Xiao-Li Zhou, Qiao-Qi Wang, Ji-Long Liu

**Author notes:** Correspondence. Ji-Long Liu.

## Abstract

Although most cells are mononuclear, the nucleus can exist in the form of binucleate or even multinucleate to respond to different physiological processes. The male accessory gland of *Drosophila* is the organ that produces semen, and its main cells are binucleate. Here we observe that CTP synthase (CTPS) forms filamentous cytoophidia in binuclear main cells, primarily located at the cell boundary. In *CTPS*^*H355A*^, a point mutation that destroys the formation of cytoophidia, we find that the nucleation mode of the main cells changes, including mononucleates and vertical distribution of binucleates. Although the overexpression of *CTPS*^*H355A*^ can restore the level of CTPS protein, it will neither form cytoophidia nor eliminate the abnormal nucleation pattern. Therefore, our data indicate that there is an unexpected functional link between the formation of cytoophidia and the maintenance of binucleation in *Drosophila* main cells.

## INTRODUCTION

Although most cells of animal organs contain a single nucleus, a considerable number of cells are multinucleate. In some cases, multinucleation is the result of cell fusion (such as placental trophoblasts or skeletal muscle) or mitosis without cytokinesis (such as liver cells or muscle cells) during tissue/cell growth [1-3]. In this case, the number of chromosome copies and nuclei in each cell increase. In other tissue/cell growth situations, such as cell division and endoduplication, each cell has only one nucleus.

Previous scientific studies show that the binucleation of hepatocytes and muscle cells occurs during the individual growth period from lactation to weaning, and the proportion of binucleated cells in stem cells gradually increases, which has been proved to be mainly determined by nutritional signals [2]. In rat muscle cells, the formation of binuclear cells mainly occurs ten days after birth[1]. However, the factors affecting binucleation remain unclear.

CTP synthase (CTPS) is an important metabolic enzyme that catalyzes the rate-limiting step of de novo synthesis of the CTP nucleotide [4]. Since 2010, it has been found that CTPS forms filamentous structures called cytoophidia in all three domains of life, including archaea, bacteria, yeast, plants, fruit flies, zebrafish and mammals [5-7].

Cytoophidia are observed in different cell types of *Drosophila*, including cells in male reproductive system [8]. It is well known that the epithelial cells of the male accessory gland (MAG) are binucleate [9]. However, it is not clear whether cytoophidia play a role in binucleation.

*Drosophila* melanogaster is a powerful tool to study human diseases, metabolism, development, innate immunity, and energy homeostasis. MAG of *Drosophila* and humans have similar tissues, anatomical structures and physiological functions [10]. The main cells of MAG of *Drosophila* provide an attractive system for studying the binuclear function of cytoophidia.

MAG is a tubular or spherical exocrine organ of internal reproductive system, of insects. Its location and function are similar to those of the mammalian prostate [11]. MAG produces and secretes components in sperm plasma, which fuse with sperm during mating and enter female copulatory sac. The epithelial cells of *Drosophila* MAG are composed of two types of cells: approximately 1000 polygonal “main cells” and 40 spherical “secondary cells”. Both types of cells have two nuclei. The binuclear system changes its apical-basal position from vertical to horizontal relative to the epithelial plane, so that the apical region of each cell has higher plasticity, thus producing a large volume of semen [12].

It is well known that substances produced by MAG play several important roles in male reproductive success. One is that the secretion of fluid provides sperm nutrition, promotes the continued development and maturation of sperm, and enhances its vitality [13]. The second role is to regulate female reproductive behavior, such as feeding, oviposition, and refusal to mate with other males, which has been elaborated by the genetically compatible insect *Drosophila melanogaster*. In the case of *Drosophila*, individuals with larger MAG volumes show higher reproductive capacity [14, 15].

MAG of *Drosophila* plays a very important role in its reproduction. We find that most of the main cells of the *Drosophila* epididymis lacking cytoophidia cannot form binucleate cells, and the presence of such mononuclear cells has no direct relationship with CTPS protein level. Our data suggest that cytoophidium formation plays a key role in the formation of binuclear cells.

## RESULTS

### CTPS forms cytoophidia in *Drosophila* male accessory glands

Previous studies have demonstrated that the reproductive systems in the adult male *Drosophila* abdomen mainly consist of four parts, namely ejaculatory duct, accessory gland, seminal vesicle, and testis (**Figure 1A**). The accessory glands composed of two symmetrical lobes (**Figure 1B**), and its epidermis has main cells and secondary cells.

**Figure 1.**
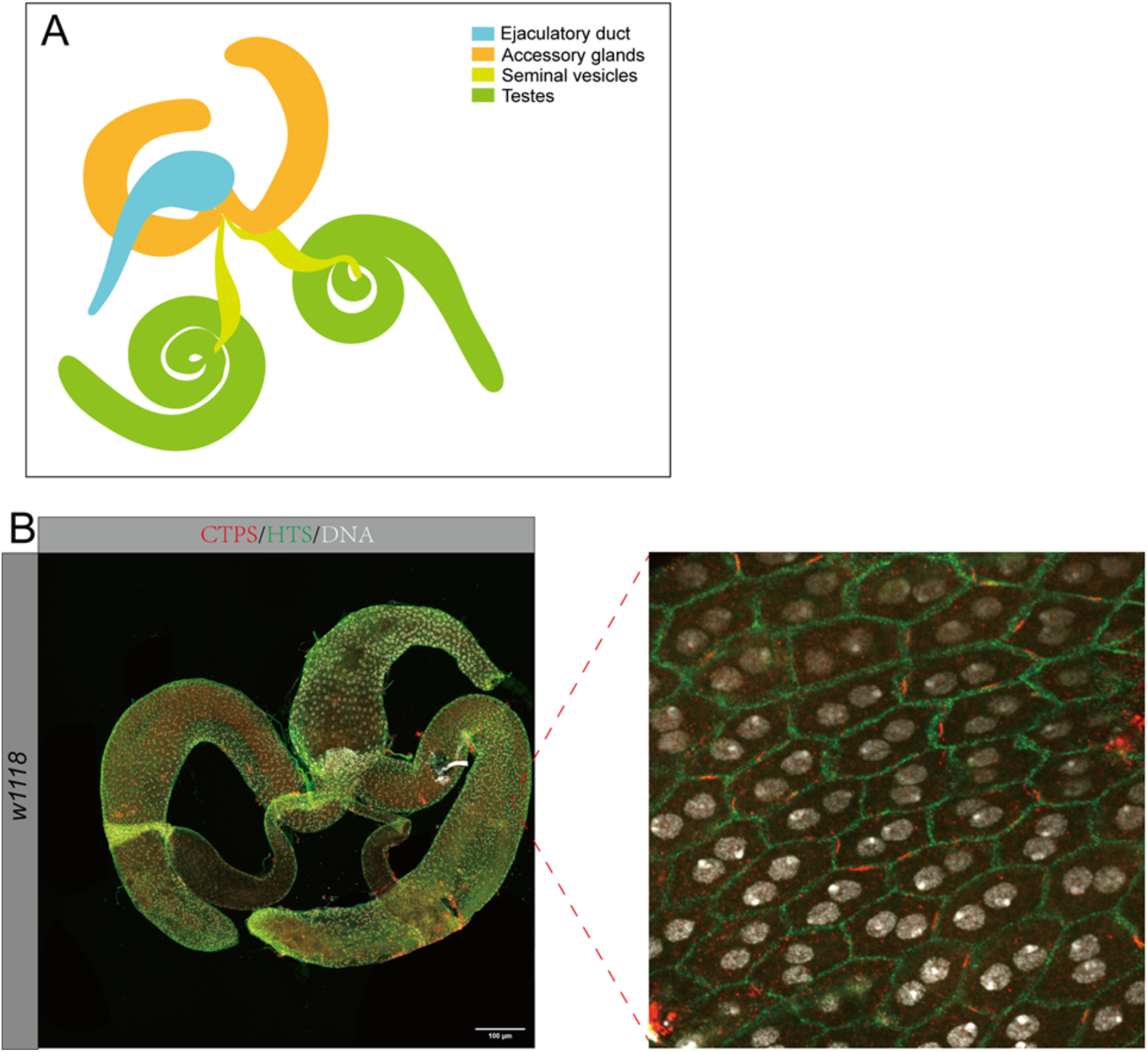
Binucleation of *Drosophila* male accessory gland cells. (A) An illustration of *Drosophila* male reproductive systems. The accessory gland is painted orange. (B) A pair of accessory glands of the *w*^*1118*^ fly, and zoom-in image on the right side. Hts (green) shows the cell boundary. CTPS (red) forms filamentous cytoophidia at the cell boundary. DNA (stained by Hoechst 33342) shows binucleation of each cell. Scale bar, 100 μm.

CTPS can form cytoophidia in multiple tissues in *Drosophila*, including the accessory glands. We first characterized polyploidy of male accessory gland epithelial cells, and the distribution of cytoophidia by immunostaining with antibodies against CTPS (labeling cytoophidia) and Hts (labeling cell membrane) and Hoechst 33342 to label DNA. We found that cytoophidia were mainly distributed in the main cells on the hexagonal cell membrane, and about two cytoophidia appear in each main cell (**Figure 2 A-F**). However, there were only a few small cytoophidia in the secondary cells (**Figure 2G**).

**Figure 2.**
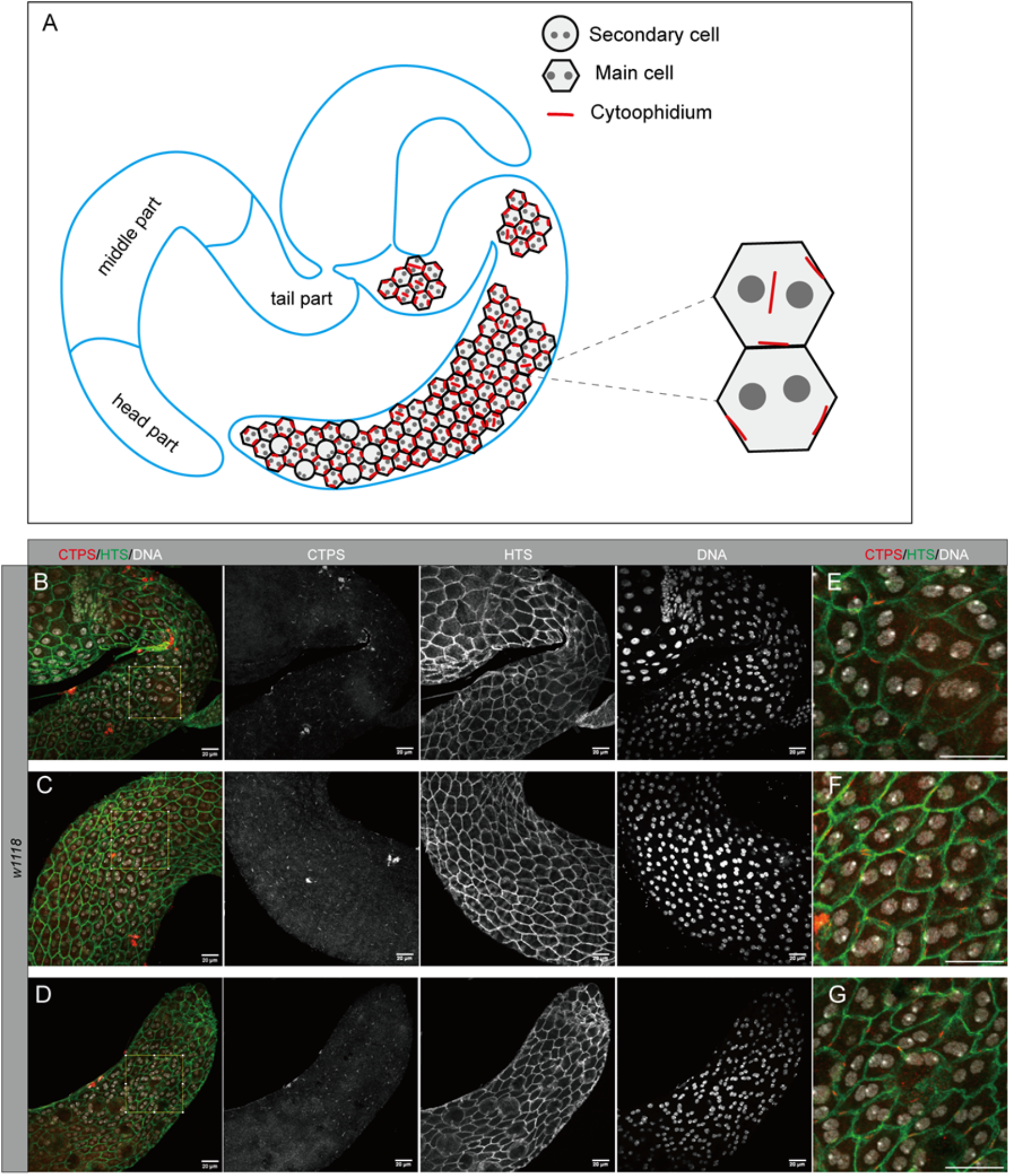
Distribution of cytoophidia in *Drosophila* accessory gland epithelia. (A) An illustration of two lobes of the accessory gland, with the distribution of cytoophidia in the main cell. (B-D) Three different parts of the accessory gland: The part near the ejaculatory duct (tail part), middle part, and the head part containing secondary cells stained by antibodies against CTPS (red), HTS (green) and by the DNA dye Hoechst 33342 (white). (E-F) Zoom-in images the tail part, middle part, and head part of the accessory gland, respectively. The fly stain is *w*^*1118*^. Scale bars, 20μm.

### *CTPS*^*H355A*^ disrupts cytoophidium formation in accessory glands

In order to further investigate the function of cytoophidia in epididymal epithelial main cells, we expressed mCherry-tagged CTPS (knock-in) in the *Drosophila* (hereinafter referred to as CTPS-mCh).

In previous studies, we knew that the 355th Histidine (H355) was an important amino acid residue in CTPS assembly to form cytoophidia. After the 355th histidine was mutated into alanine in vitro, CTPS could not be assembled (**Figures 3A and B**). Then, we expressed histidine to alanine (knock-in) at the 355th position of CTPS mutation in CTPS mCh *Drosophila* (hereinafter referred to as H355A).

**Figure 3.**
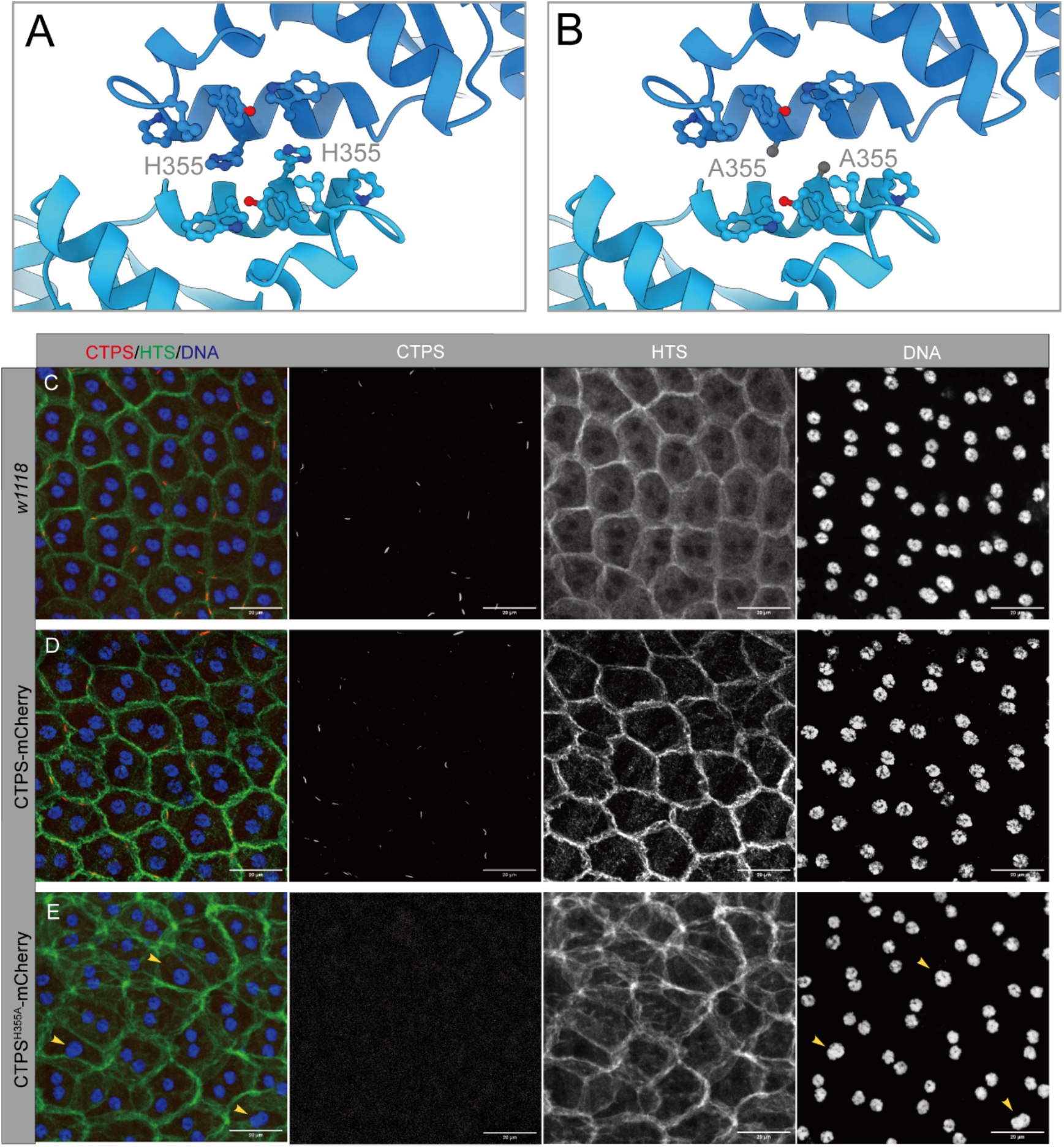
*CTPS*^*H355A*^ disrupts cytoophidium formation in accessory glands. (A) Structure of CTPS shows that H355 lies at the tetramer-tetramer interface. (B) H355A mutation of CTPS weakens the tetramer-tetramer interaction. (C) In *w*^*1118*^ fly, staining by an antibody against CTPS (red in the merge image) shows that cytoophidia lie at the cell boundary (HTS, green in the merge image). Each cell contains two nuclei (DNA, stained by Hoechst 33342, blue in the merge image). white). (D) In *CTPS-mCherry* transgenic fly, CTPS-mCheery (red in the merge image) shows that cytoophidia lie at the cell boundary (HTS, green in the merge image). Each cell contains two nuclei (DNA, stained by Hoechst 33342, blue in the merge image). (E) In *CTPS*^*H355A*^*-mCherry* transgenic fly, CTPS^H355A^-mCheery (red in the merge image) shows no detectable cytoophidia. The cell boundary is labelled by HTS (green in the merge image). Some cells (yellow arrowheads) only contain one nucleus (DNA, stained by Hoechst 33342, blue in the merge image). Scale bars, 20 μm.

To directly test the function of cytoophidia in the main cells of accessory glands, we took CTPS-mCh *Drosophila* as the wide-type and H355A mutant as the test object. By comparing with *w*^*1118*^, we found that it did not affect CTPS assembly in main cells (**Figure 3C and D**).

To observe the difference between the main cells in the accessory gland of H355A mutant and the wild-type, we used an antibody against CTPS to label CTPS protein, anti-HTS on the cell membrane to label HTS, and the DNA dye Hoechst 33342 to label the nucleus. We found that CTPS-forming cytoophidia were mainly distributed on the hexagonal cell membrane labeled with HTS in the main cell (**Figure 3C and D**), and CTPS could not be assembled in the H355A mutant (**Figure 3E**).

### Mononucleation occurs in *CTPS*^*H355A*^ main cells

Secondary cells are mainly distributed in the head of accessory glands. Each secondary cell is surrounded by five or six main cells. In the wild-type control, each secondary cell was slightly larger than the main cell (**Figure 4A**). In the head of *CTPS*^*H355A*^ accessory glands, the secondary cells seemed to be much larger than the main cells (**Figure 4B**).

**Figure 4.**
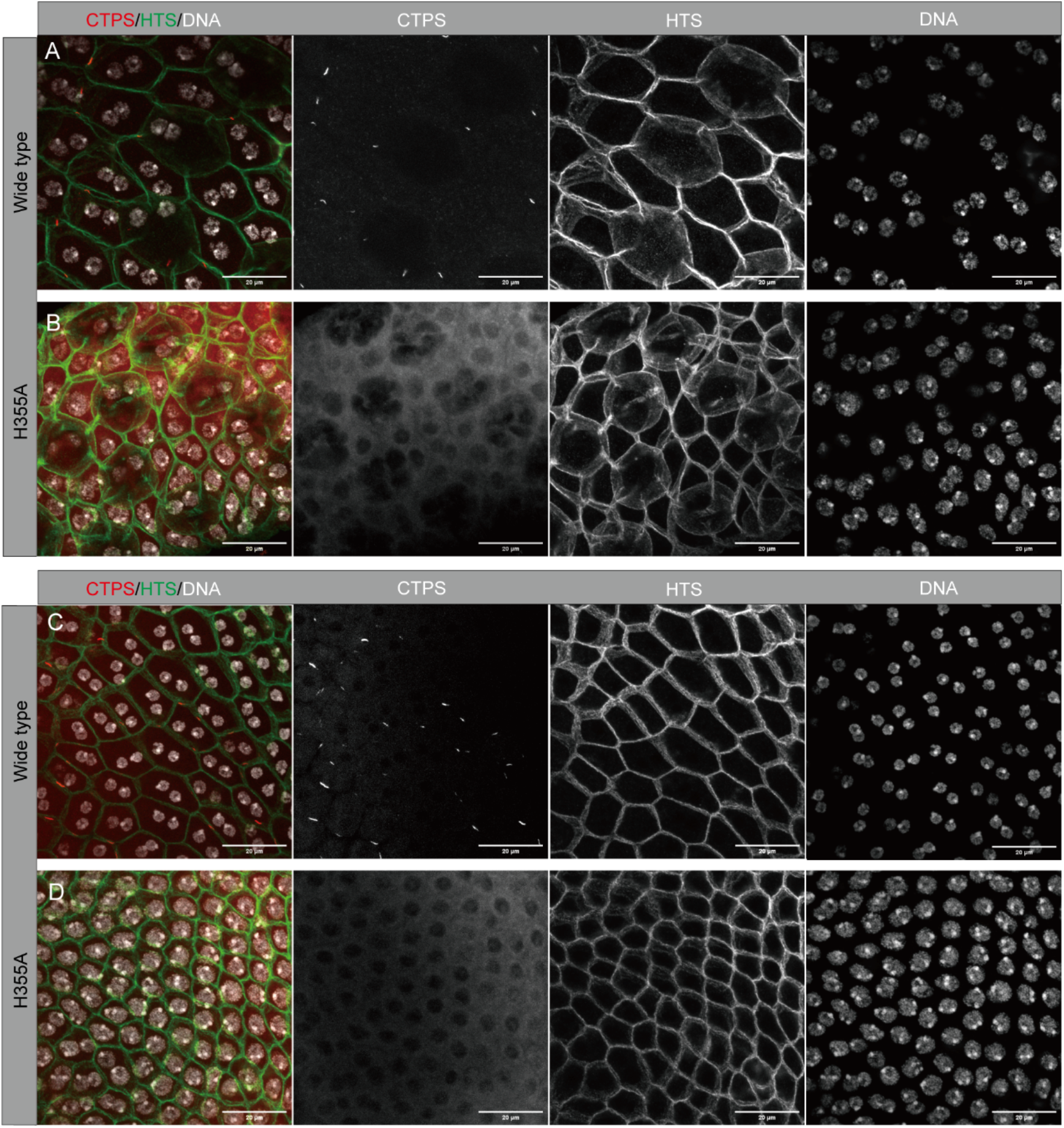
Mononucleation increases in *CTPS*^*H355A*^ main cells. (A, B) The head part of the accessory gland contains both main cells and secondary cells. A is wide-type control, and B is *CTPS*^*H355A*^ mutant. Cytoophidia appear in wild-type control but not in the *CTPS*^*H355A*^ mutant. Note that the ratio between the average area of main cells and that of secondary cells appears smaller in *CTPS*^*H355A*^ mutant than that in wild-type control. (C, D) The middle part of the accessory gland contains main cells only. A is wide-type control, and B is *CTPS*^*H355A*^ mutant. Cytoophidia appear in wild-type control but not in *CTPS*^*H355A*^ mutant. Note that the average area of main cells appears smaller in *CTPS*^*H355A*^ mutant than that in wild-type control. (D) In this image, the most majority cells contain only one nucleus, in contrast to binuclei in all cells shown in wild-type (C). In the merge images of A-D, CTPS (red), HTS (green), and DNA (white). Scale bars, 20 μm.

The middle of accessory gland contains main cells, but no secondary cells. In the wild-type accessory glands, CTPS formed obvious cytoophidia in the main cells, each of which contains two nuclei (**Figure 4C**). In the middle of *CTPS*^*H355A*^ accessory gland, CTPS did not form cytoophidia. Interestingly, we observed that some main cells had only one nucleus (**Figure 4D**). The proportion of mononucleation could be different among individual mutant flies. In some cases, almost all main cells contained only one nucleus (**Figure 4D**).

The cell size of the main cells of the *CTPS*^*H355A*^ mutant seemed to be much smaller than that of the wild-type control (**Figure 4C and D**). However, the nuclear size of the monoclear main cells of *CTPS*^*H355A*^ mutant was significantly larger than that in the wild-type control (**Figure 4C and D**).

### Vertical distribution of binuclei exists in *CTPS*^*H355A*^ main cells

In the wild-type accessory glands, each main cell contains two nuclei, which are horizontally distributed in the epithelium. The distance between the the two nuclei is relatively large. The horizontal distribution of binuclei improves the ductility and mobility of cells to a certain extent, which is more conducive to the secretion of a large amount of semen.

Three types of nucleation were observed in *CTPS*^*H355A*^ main cells: 1) two nuclei arranged horizontally, similar to the wild-type control; 2) mononucleus, as described above; and 3) two nuclei arranged vertically to the epithelium (**Figure 5A-F**).

**Figure 5.**
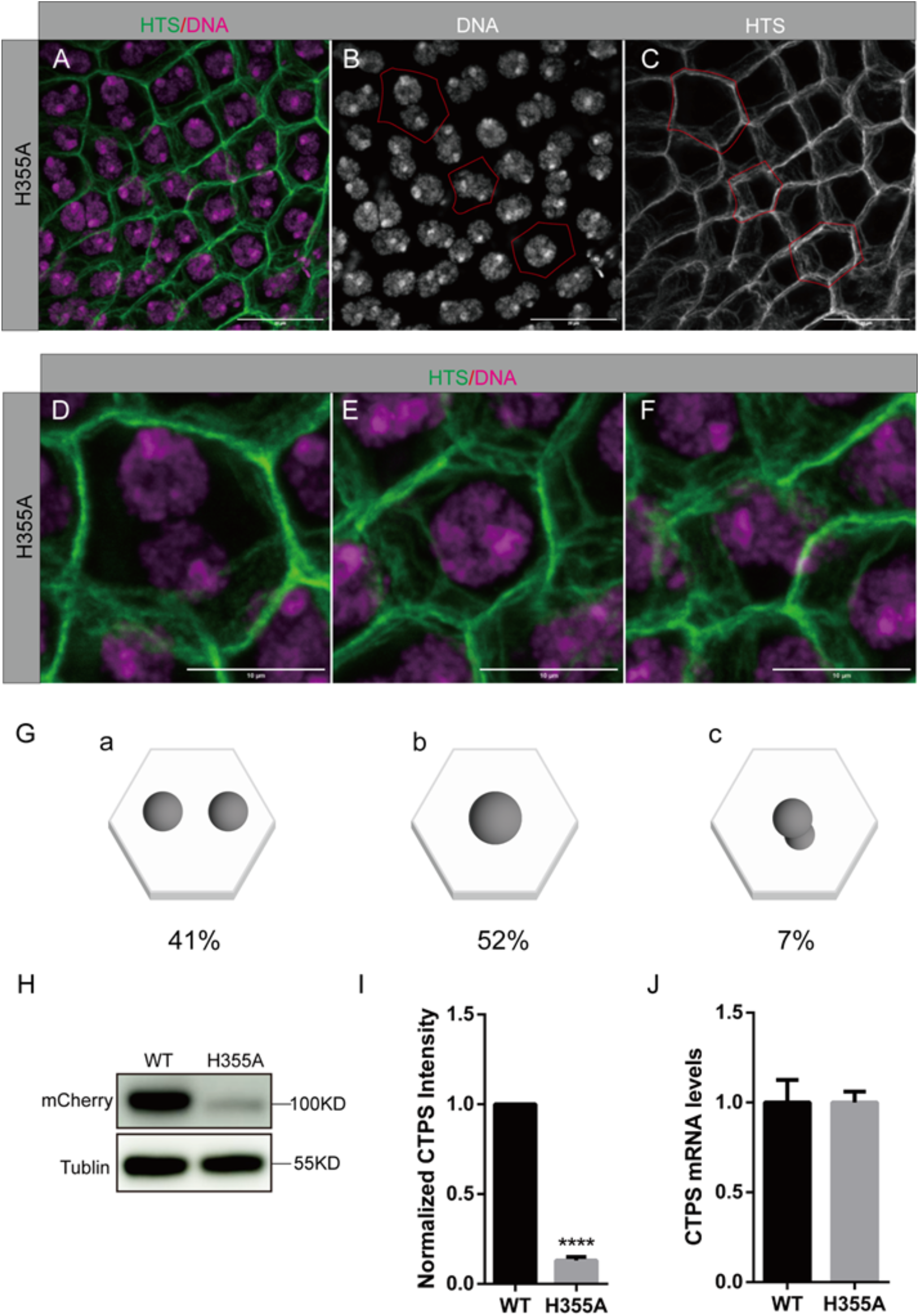
Three nucleation types in *CTPS*^*H355A*^ main cells. (A-C) In *CTPS*^*H355A*^ mutant, there are three nucleation types of main cells. Hts (green in A, white in C) shows the cell boundary. DNA (stained by Hoechst 33342, mageta in A, white in B) shows nucleation of each cell. Red lines in B and C outline cells with three nucleation types (Zoom-in images are shown in D-F). Note the differences in cell size and morphology. Scale bars in A-C, 20 μm. (D-F) Three nucleation types of main cells in *CTPS*^*H355A*^ mutant: normal binucleated main cell (D), mononucleated main cell (E), and main cell with two nuclei arranged vertically (F). Scale bars in D-F, 20μm. (G) The proportion of the three nucleation types of main cells: 41% for normal cell with two nuclei arranged horizontally to the epithelium (a), 52% for mononucleated main cell (b), and 7% for main cell with two nuclei arranged vertically to the epithelium (c). (H) Western blot detected by anti-mCherry and anti-tubulin antisera on lysates of male reproductive systems from *wild-type* and *CTPS*^*H355A*^ mutant flies. (I) Quantification analyses of the western blot of CTPS. (J) Real-time PCR analysis of CTPS transcriptional levels in the male reproductive systems from *wild-type* and *CTPS*^*H355A*^ mutant flies. ns, non-significant; ****, p<0.0001; Mann-Whitney test.

In *CTPS*^*H355A*^ accessory glands, the size of main cells with vertically distributed nuclei was smaller than that of main cells with horizontally distributed nuclei. The distance between the two nuclei arranged vertically seemed to be much smaller than that between the nuclei arranged horizontally (**Figure 5D and F**). Next, we quantified the proportion of three types of nucleation in *CTPS*^*H355A*^ main cells. 41% of *CTPS*^*H355A*^ main cells showed two nuclei arranged horizontally, 52% mutant main cells only contained only one nucleus, and 7% cells had with two nuclei arranged vertically (**Figure 5G**).

### CTPS protein level decreases in *CTPS*^*H355A*^ mutant

To explore whether the CTPS content in the wide-type and H355A is different, we used western blotting and qPCR to detect the CTPS protein and mRNA content respectively. Western blotting showed that the protein content of CTPS in H355A was significantly lower than that of wide-type (**Figure 5H and I**). However, there was no significant difference in CTPS mRNA (**Figure 5J**).

### Mononucleation occurs in main cells overexpressing *CTPS*^*H355A*^

To eliminate the influence of protein content, we overexpressed CTPS protein and H355A mutated protein in the main cells of the accessory gland through *Drosophila* UAS-Gal4 transgenic system. We constructed Actin-Gal4 driving UAS-CTPS mCherry in wide-type and the H355A mutant (hereinafter referred to as *Actin>CTPS-mCh* and *Actin>H355A*, respectively).

Overexpression of CTPS led to the formation of large and long cytoophidia, which were distributed at the edge of the hexagonal cell membrane of the main cells (**Figure 6A-H**). There were come cytoophidia between the two nuclei of the main cell, which seemed to separate the nuclei.

**Figure 6.**
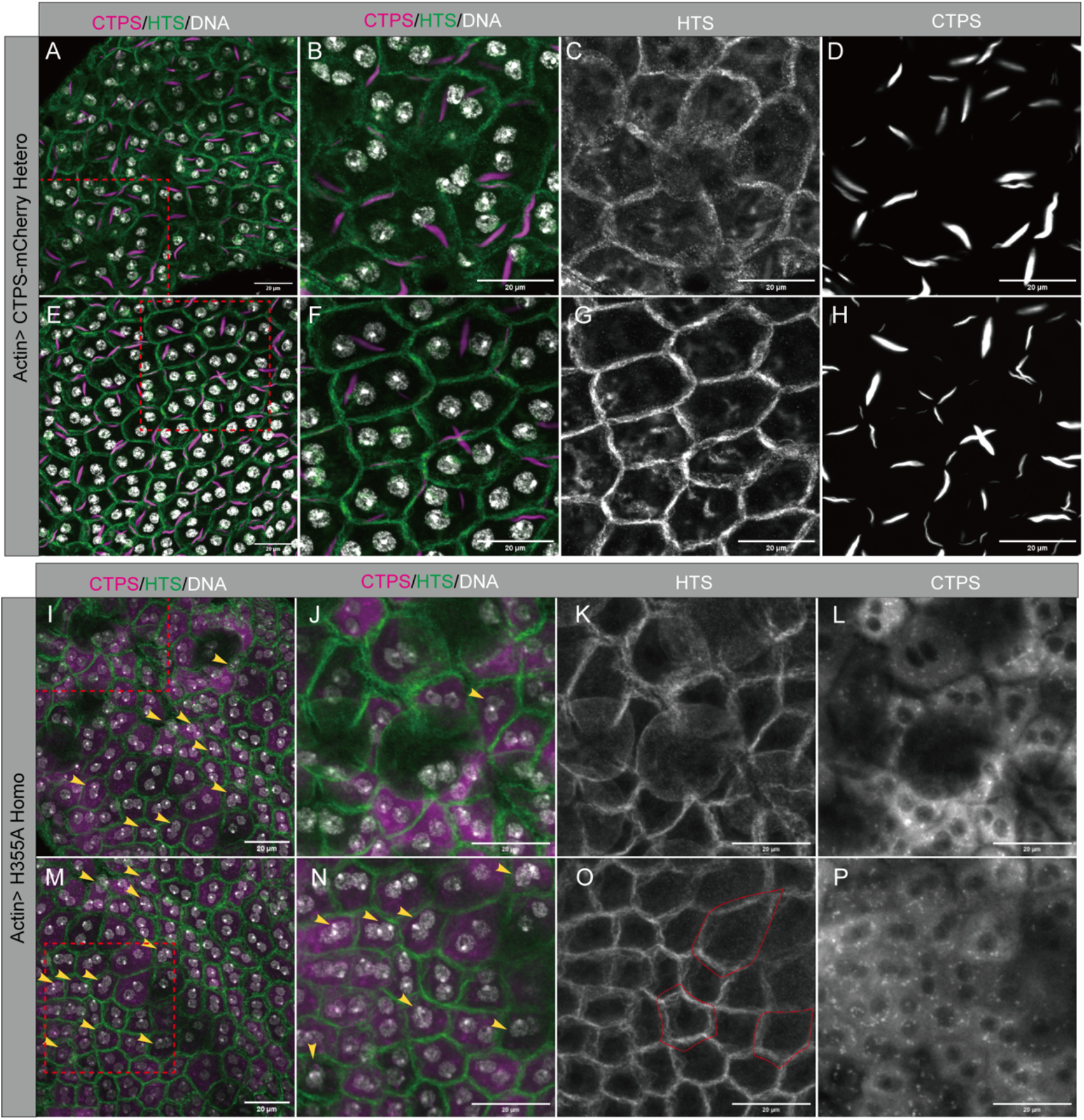
Mononuceation occurs in overexpressed *CTPS*^*H355A*^ main cells. (A-H) Heterozygous accessory gland overexpressing CTPS (driven by *Actin-Gal4*). The head part (A, zoom-in image in B) and middle part (E, zoom-in image in F) of the accessory gland. Note that overexpressed CTPS forms giant cytoophidia (mageta in A, B, E, F; white in D, H) near the cell boundary (Hts, green in A, B, E, F; white in C, G). In some cells (yellow arrows), a giant cytoophidium lies at the central space separating the two nuclei (DNA, stained by Hoechst 33342, white in A, B, E, F). (I-P) Homozygous accessory gland overexpressing *CTPS*^*H355A*^ (driven by *Actin-Gal4*). The head part (I, zoom-in image in J) and middle part (M, zoom-in image in N) of the accessory gland. Note that overexpressed *CTPS*^*H355A*^ shows increased diffused CTPS signal (mageta in I, J, M, N; white in L, P) but no detectable cytoophidia. Cell boundary is labeled by Hts (green in I, J, M, N; white in K, O). Some main cells (yellow arrowheads) show abnormal nucleation (either containing only one nucleus or having two nuclei arranged vertically). Red lines in O outline abnormal cell shapes. Scale bars, 20 μm.

In *CTPS-mCherry*^*H355A*^ accessory glands, CTPS showed an increased but diffused distribution pattern. Mononucleate cells were observed in the *CTPS-mCherry*^*H355A*^ flies driven by *Actin-Gal4*, while no mononucleation was found in the control flies (*Actin>CTPS-mCh*) (**Figure 6I-O**). In *CTPS-mCherry*^*H355A*^ accessory glands, the cell size of mononucleate main cells seemed to be smaller than that of binucleate main cells.

Western Blotting analysis of male reproductive system showed that the CTPS protein level of heterozygous *actin>CTPS-mCh* was similar to that in homozygous *actin>H355A* flies. However, the CTPS protein level of the homozygote *actin>CTPS-mCh* was twice that of the homozygous *actin>H355A* (**Figure 7A-B**).

**Figure 7.**
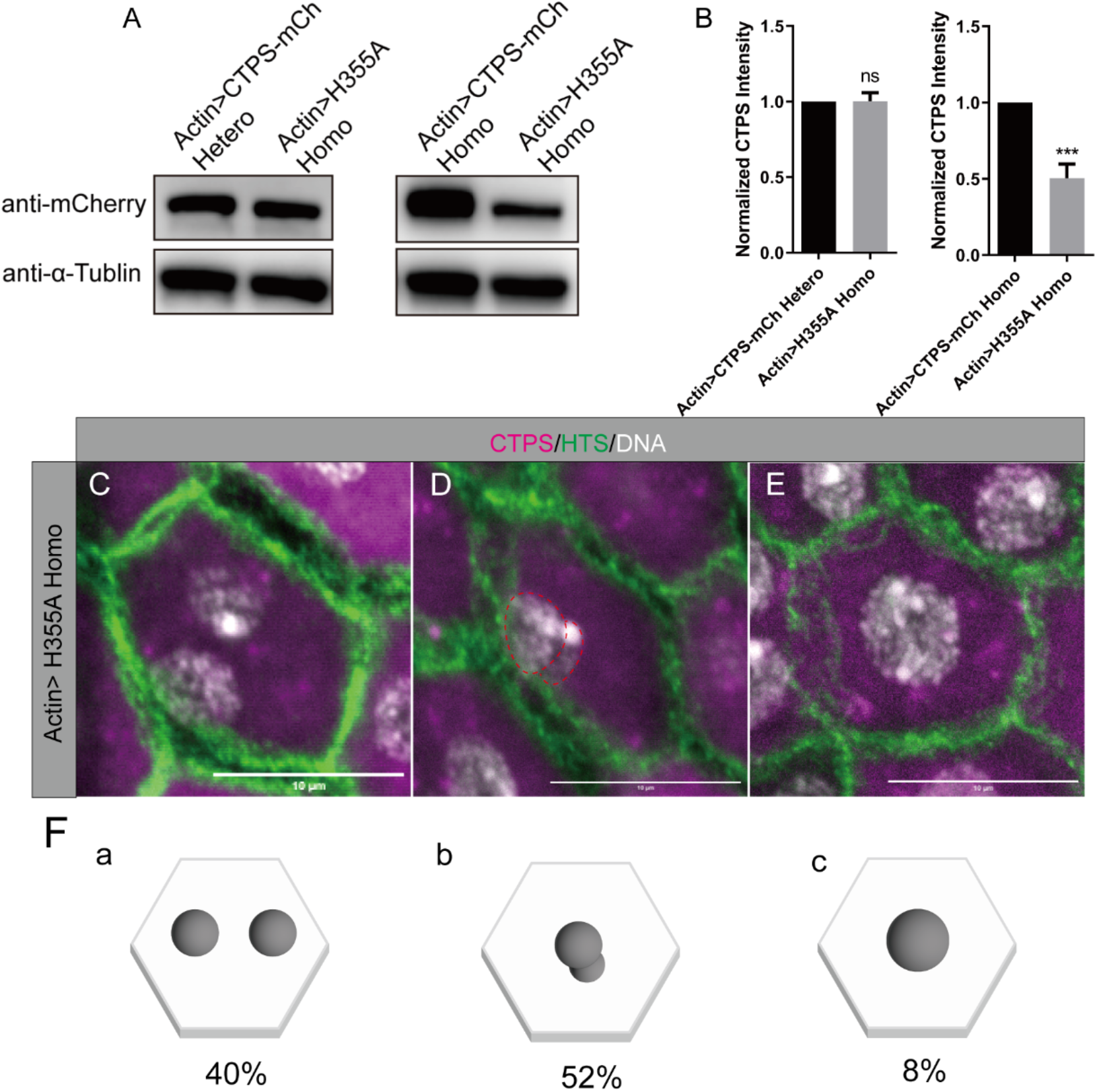
Three nucleation types in main cells overexpressing *CTPS*^*H355A*^. (A) Western blotting, detected by antibodies against mCherry and tubulin, on lysates of male reproductive systems from flies overexpressing *wild-type CTPS-mCherry* and *CTPS*^*H355A*^*-mCherry* (both driven by *Actin-Gla4*), respectively. The genotypes are 1) Actin>CTPS-mCh Hetero; 2) Actin>H355A Homo; 3) Actin>CTPS-mCh Homo; 4) Actin>H355A Homo. (B) Quantification of Western blotting with the genotypes shown in A. **, p<0.01, Mann-Whitney test. (C-E) Three nucleation types of main cells overexpressing *CTPS*^*H355A*^ (driven by *Actin-Gal4*): normal binucleated main cell (C), main cell with two nuclei arranged vertically (D), and mononucleated main cell (E). Scale bars in C-E, 10 μm. (F) The proportion of the three nucleation types of main cells overexpressing *CTPS*^*H355A*^ (driven by *Actin-Gal4*): 40% for normal cell with two nuclei arranged horizontally to the epithelium (a), 52% for main cell with two nuclei arranged vertically to the epithelium (b), and 8% for mononucleated main cell (c).

### Overexpressing *CTPS*^*H355A*^ leads to an increase in binuclear vertical distribution

According to the nucleation types in the *Actin>H355A* mutant, we found that the proportion of horizonally binucleated cells was 40%, the proportion of cells with vertically distributed nuclei was 52%, and the proportion of mononuclear cells was 8% (**Figure 7C-F**).

Although compared with H355A mutant without overexpression of CTPS, the proportion of mononuclear main cells was significantly reduced, there were still a considerable number of mononuclear main cells, indicating that cytoophidia affected the nuclear division of the main cells to some extent.

Compared with H355A mutants, the CTPS protein content of CTPS overexpression mutants increased, and the number of main cells with vertically distributed nuclei also increased from 7% to 52%. The nuclei in the vertically distributed main cells were smaller than those of the mononuclear main cells (**Figure 5D-G; Figure 7C-F**).

In H355A mutants, the main cells with vertically distributed nuclei appeared only when the cell size shrank. However, in CTPS overexpression mutants, we found a large number of cells with vertically distributed nuclei, even in normal-sized main cells. In the overexpressed wild-type main cells, there was almost no nuclear vertical distribution of the main cells (**Figure 7C-F**).

## DISCUSSION

Here, we observe the distribution of cytoophidia in male accessory glands of *Drosophila melanogaster*. In *CTPS*^*H355A*^ mutant main cells, both cytoophidium formation and binucleation are impaired. In addition, overexpression of *CTPS*^*H355A*^ can prevent cytoophidium formation and affect the binucleation of main cells. Our results indicate that cytoophidia safeguard binucleation of the main cells in *Drosophila* male accessory glands.

Although most eukaryotes are diploid, there are still many cells with more than two sets of chromosomes. In some mammalian tissues and organs, a large number of tetraploid cells will be produced in the physiological or pathological processes [16-18]. Interestingly, phenotypic analysis shows that increasing ploidy can make cellular physiology change to a specific functional direction. However, the production of polyploid cells can also trigger cell transformation and tumor formation [19]. It has been clearly shown that liver proliferation is associated with ploidy during development and in challenging situations, such as partial hepatectomy or after oxidative damage [20-22]. In this context, determining which factors trigger the production of polyploid cells may bring new insights into the physiology of functions of these cells.

In previous studies, we found that cytoophidia exist in a variety of organisms and cell types, and a series of studies found that cytoophidia play an important role in the structure and function of cells [23, 24]. Here we observe cytoophidia in the main cells of *Drosophila* male accessory glands with binuclear system. The connection between cytoophidia and the binuclear system is explored in the *CTPS*^*H355A*^ mutant, in which CTPS is unable to form cytoophidia.

We are surprised to find that many main cells in the *CTPS*^*H355A*^ mutant cannot form a binuclear system. Compared with the horizontal distribution of the binuclear system in the wild-type main cells, many vertically distributed binuclear cells appear in the mutant. The horizontal distribution of the binuclear system in the main cells of *Drosophila* has important physiological significance for the production of semen. The effect of the absence of the binuclear system on the accessory gland function of *Drosophila* needs further studies.

In the *CTPS*^*H355A*^ mutant, we find that although CTPS is the same as the wild-type control flies at the transcription level, its protein content is significantly reduced. Eliminating the factor of the protein content, our experimental results show that the main cells in which CTPS fails to assemble into cytoophidia can still leads to mononucleation. The number of vertically distributed cells overexpressing the H355A mutant is significantly increased compared with the wild-type main cells. Both monocytes and the main cells with vertically distributed binuclei are smaller than those with horizontal binuclei. Moreover, we also find that, in the control *Drosophila* accessory glands, the main cells are mainly distributed on the hexagonal cell membrane, so we speculate that the formation of cytoophidia may also have a certain impact on the cell size.

In conclusion, our results suggest that cytoophida are crucial for the binucleation of the main cell of *Drosophila* accessory glands. Therefore, our study provides the potential function of cytoophidia in the regulation of cell division and development.

## MATERIALS AND METHODS

### Fly strains and genetics

Fly stocks were raised on standard cornmeal food at 25°C. The following strains were used: *w*^*1118*^, *C-terminal mChe-4V5 tagged CTPS* knock-in fly produced in our laboratory, *H335A mutated with mChe-4V5 tagged CTPS* Knock-in fly, *Actin-Gal4, Sp/Cyo*; *Sb/Tm6B*, heterozygote of *UAS CTPS-mCh-OE*, homozygote of *UAS CTPS-mCh-OE*, and homozygote of *UAS CTPS*^*H355A*^*-mCh-OE*.

The *actin-Gal4; UAS CTPS-mCh-OE* strain was generated by crossing the homozygous *UAS CTPS* and *Actin-Gal4* fly strains with the double balancer flies for one generation. The target generation were crossed for two generations to obtain the homozygous and heterozygous *actin-Gal4; UAS-CTPS-mCh-OE* flies. Homozygous *actin-Gal4; UAS-CTPS*^*H355A*^*-mCh-OE* flies were produced in a similar way.

### Immunohistochemistry

The tissues of male *Drosophila* for 3-5 days were dissected from at least three technical replicates, and each replicate contained more than 5 animals. Flies were dissected in Grace’s Insect Medium (Cat. No. 11605045; Invitrogen, Carlsbad, California, USA), and then fixed in 4% formaldehyde (Cat. No. F8775; Sigma, Darmstadt, Germany) in phosphate-buffered saline (PBS) for 10 min, followed by washing in PST (0.5% horse serum +0.3% Triton X-100 in PBS). The samples were incubated overnight in primary antibodies at room temperature. They were then washed briefly with PST and incubated in secondary antibodies and Hoechst 33342 (1:100,000; Thermo Fisher, Rockford, IL, USA) for DNA staining overnight at room temperature. Finally, the sample was mounted on the slide for confocal microscopy.

Primary antibodies used in this study are as follows: rabbit anti-CTP synthase 1/2 (y88) (1:1000, Santa Cruz Biotechnology, Cat. No. sc-134457) and mouse anti-HTS (1:1000, Developmental Studies Hybridoma Bank, Cat. No. AB_528070). Secondary antibodies used in this study are as follows: donkey anti-rabbit IgG, Cy5 (1:1000; Jackson ImmunoResearch Laboratories, West Baltimore Pike, West Grove, PA); and goat anti-mouse IgG, Alexa Fluor 488 (1:1000; Cat. No. A28175; Thermo Fisher, Waltham, MA, USA).

### Microscopy

Images were ftaken under 40 X or 63 X objectives on a laser-scanning confocal microscope (Zeiss LSM980; Leica microsystems; Germany). Images were analyzed using ImageJ (version 1.43 U; National Institutes of Health, Bethesda, Maryland, USA). The most representative images were displayed with maximum intensity projection.

### Cell counting

The number of cells was counted using ImageJ. At least three biological replicates were quantified.

### Quantitative RT-PCR

Total RNA was collected from male reproductive system of adult *Drosophila* by RNA preparation kit (TransGen, Beijing, China). RT-PCR kit (Takara, Tokyo, Japan) was used to construct reverse transcription to collect cDNA. The cDNA was used for real-time PCR using SYBR Green master PCR Mix (Vazyme, Nanjing, China) in triplicate. The raw data were collected on the Quantstudio 7 (Life Technologies) detection system. The transcript level was normalized to rp49. At least three biological replicates were quantified. The Primers used are shown as follows: CTPS, forward primer: 5’-CCGAGGTGGATCTGGATCT-3’, reverse primer: 5’-GTGGCTTTGAAGATCCCTGA-3’; rp49, forward primer: 5’-GCTAAGCTGTCGCACAAA-3’, reverse primer: 5’-TCCGGTGGGCAGCATGTG-3’.

### Western blotting

The adult male testes of *Drosophila melanogaster* were collected into the lysis buffer RIPA (Meilunbio, Dalian, China), combined with the protease inhibitor cocktail (Bimake, Shanghai, China) for Western blot, and then ground with 1 mm Zirconia beads in Sonicator (Shanghai Jing Xin, Shanghai, China). After 30 min, the samples were lysed on ice. At 10000 g at 4°C, the samples were centrifuged for 10 min. The 1X protein loading buffer was pipetted to the supernatant and boiled at 95°C for 10 min. Subsequently, the samples were run with 10% SDS-PAGE gel and transferred to PVDF membrane. At room temperature, the membrane was incubated with 5% w/v nonfat dry milk dissolved with 1X TBST for 1 h.

Then the sample was incubated with primary antibodies in 5% w/v nonfat milk at 4°C with gentle shaking overnight. The following primary antibodies were used in this study: Monoclonal antibodies against mCherry Tag (Cat. No. A02080, Abbkine, Beijing China) and mouse anti-α-Tubulin antibody (Cat. No. T6199, Sigma). The membrane was washed with shaking three times each time for 5 min, then incubated with secondary antibodies (anti-mouse IgG, HRP-linked antibody, Cell Signaling, Danvers, MA, USA) diluted with 5% w/v nonfat milk at room temperature for 1 h.

Amersham Imager 600 (General Electric, Boston, MA, USA) and Pierce ECL Reagent Kit (Cat. No. 32106, Thermo Fisher, USA) was constructed for chemiluminescence immunoassay. Protein levels were quantified on ImageJ and normalized to tubulin. At least three biological replicates were quantified.

## ACKNOWDGMENTS

We thank the Molecular Imaging Core Facility (MICF) at School of Life Science and Technology, ShanghaiTech University for providing technical support. We also thank Professor Clive Wilson for helpful discussion. This research was funded by Ministry of Science and Technology of China (grant number 2021YFA0804700), National Natural Science Foundation of China (grant number 31771490), Shanghai Science and Technology Commission (grant number 20JC1410500) and the UK Medical Research Council (grant numbers MC_UU_12021/3 and MC_U137788471).

